# Photoacoustics-guided Real-Time Closed-loop Control of Magnetic Microrobots through Deep Learning

**DOI:** 10.1101/2025.08.13.670011

**Authors:** Richard Nauber, Johanna Hoppe, David Castellanos Robles, Mariana Medina-Sánchez

**Author notes:** Corresponding author: M. Medina-Sánchez.

## Abstract

Medical microrobots promise to increase the efficacy and reduce the invasiveness of certain medical procedures in the future. Real-time tracking of the microrobot, actuation, and closed-loop control of its position under *in vivo* conditions is crucial to fulfill the task at hand.

We present a system for closed-loop control of magnetic microrobots using dual-mode ultrasound and photoacoustic imaging. It employs GPU-accelerated beamforming and tracking to achieve real-time operation with a closed-loop cycle time of 100 ms. Artifacts from simultaneous imaging and magnetic actuation are suppressed through time-multiplexing.

To address the challenge of detecting microrobots in low-contrast, strong-background images, we implemented real-time Deep Learning-based tracking. A custom dataset of various types of microrobots is curated from long-duration closed-loop control measurements and employed to fine-tune a pre-trained detection model.

We introduce a platform for real-time closed-loop control of microrobots and demonstrate its performance with a 300 μm spiral-shaped microrobot following a figure-of-8 shape under photoacoustic imaging guidance. The localization error is evaluated against an optical reference measurement. Our results show that photoacoustic-based tracking significantly outperforms ultrasound tracking, with the deep learning approach further reducing missed detections. This demonstrates the algorithm’s ability to generalize to a previously unseen type of microrobot. We envision this platform to advance medical microrobotics research by providing real-time closed-loop control of untethered microrobots under deep tissue.

## I. Introduction

The emerging field of medical microrobotics offers the vision to reduce the invasiveness of some medical procedures and improve their outcome. For instance, in reproductive medicine, directly transferring an embryo with a microrobotic carrier to the fallopian tube after *in vitro* fertilization may enhance the implantation success rate and replace an invasive laparoscopic intervention [1]. However, for performing *in vivo* tasks with untethered microrobots, it is essential to have maneuverability and real-time trackability under deep tissue [2]. Magnetic microrobots, which are remote-controlled through external magnetic fields and are tracked with ultrasound (US) or photoacoustics (PA) imaging, can provide that under clinically relevant constraints [3]. They use non-ionizing radiation and are considered low-cost and safe in general [4].

We present a flexible closed-loop position control system for magnetic microrobots based on multi-modal imaging feedback. It is tailored to *in vivo* applications and supports dual-mode PA and US imaging while simultaneously actuating the microrobots with a magnetic field. Imaging artifacts, which emerge from an electromagnetic coupling of the strong pulsed actuation currents with the low-voltage ultrasound signals, are addressed through a precise synchronization and time-multiplexing of imaging and actuation. The challenge of localizing microrobots in complex environments through low signal-to-noise-ratio images is addressed with a deep-learning (DL) based detection. It can be used instead of the standard threshold-based detection to increase the robustness under these conditions by leveraging a learned representation of the microrobots in the images. Real-time performance is achieved by utilizing GPU acceleration for both detection methods.

We demonstrate the closed-loop control of a 300 μm spiral-shaped microrobot with PA-based feedback and evaluate the localization performance by comparison with optical reference measurements. Based on the obtained data, different variants of detection can be evaluated objectively.

## II. Materials and Methods

### A. Real-time imaging and actuation

The closed-loop system comprises of tightly coupled imaging and actuation components (Fig. 1). Ultrasound emission and data acquisition are performed with an open ultrasound research platform (us4R, us4us, Warsaw, Poland), which allows real-time raw-data access through an open-source software [5]. It is connected to a 256 elements CMUT probe (L22-8 Kolo Medical) with a center frequency of 17 MHz. The excitation for PA imaging is provided through a pulsed laser (Q-TUNE-E10-SH, Quantum Light Instruments Ltd., Lithuania). The raw US and PA data is beamformed on a GPU (RTX2080, Nvidia, Santa Clara, USA) and the threshold-based localization using CuPy [8] or the DL-based localization (cf. Section II.C) is performed. This is necessary to meet the overall timing budget for closed-loop control with 10 Hz sample rate (Fig. 2). Afterwards, on the CPU, the linking of the detected objects between frames is performed using TrackPy [10]. Then the deviation from the planned trajectory the respective magnetic field vector and coil currents for the next time step are calculated. A custom orchestration module based on an RP2040 microcontroller (Raspberry Pi Ltd., Cambridge, UK) with an 8-channel H-bridge driver interfaces with a magnetic field generator (MFG-100-i, field enhancement module 10 mm, Magnebotix AG, Switzerland).

**Figure 1:**
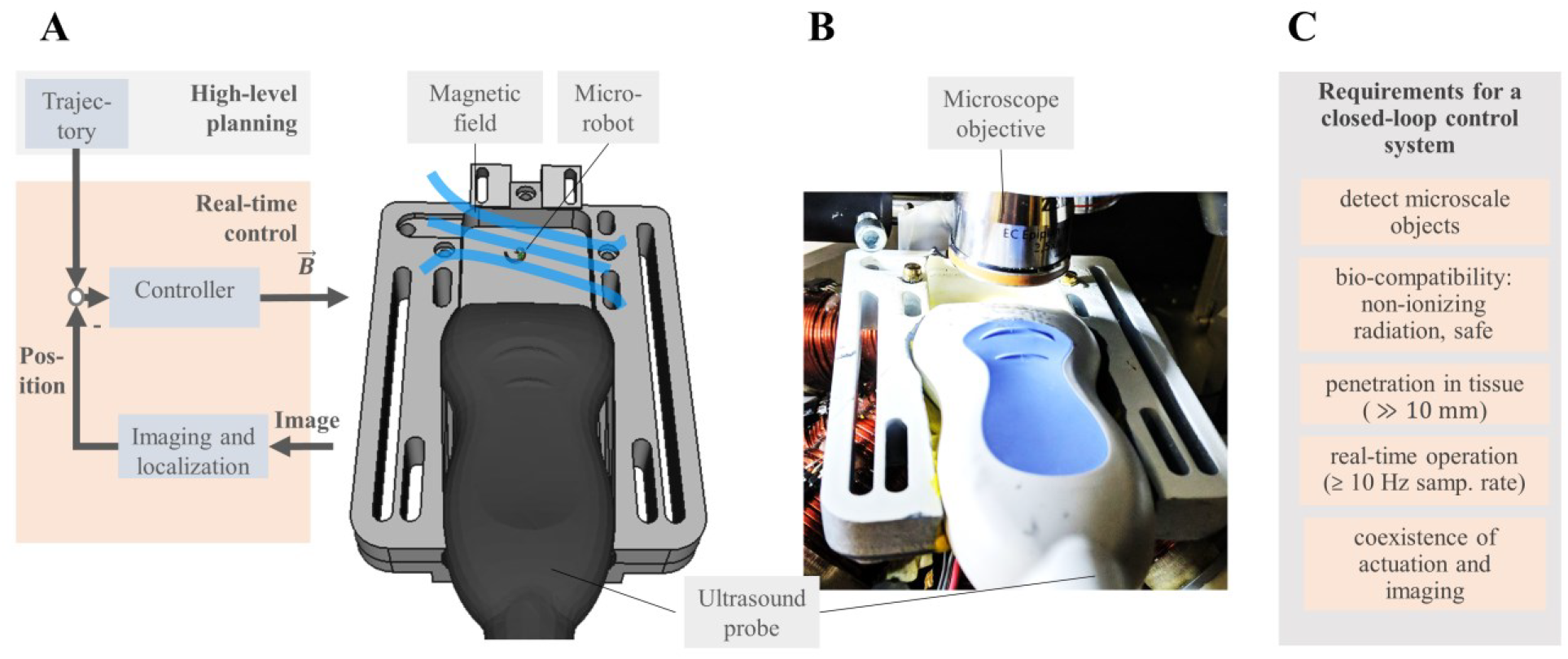
(A) High-level overview and experimental setup for evaluating closed-loop control system’s performance on magnetic microrobots. (B) An US array images a plane above a computer-controlled magnetic field generator, where a microrobot can be actuated. Pulsed laser illumination allows for PA imaging as well. Auxiliary optical imaging can be performed though a microscope from the top at the same time. (C) General requirements for a closed-loop motion control system in context of medical microrobotics.

**Figure 2:**
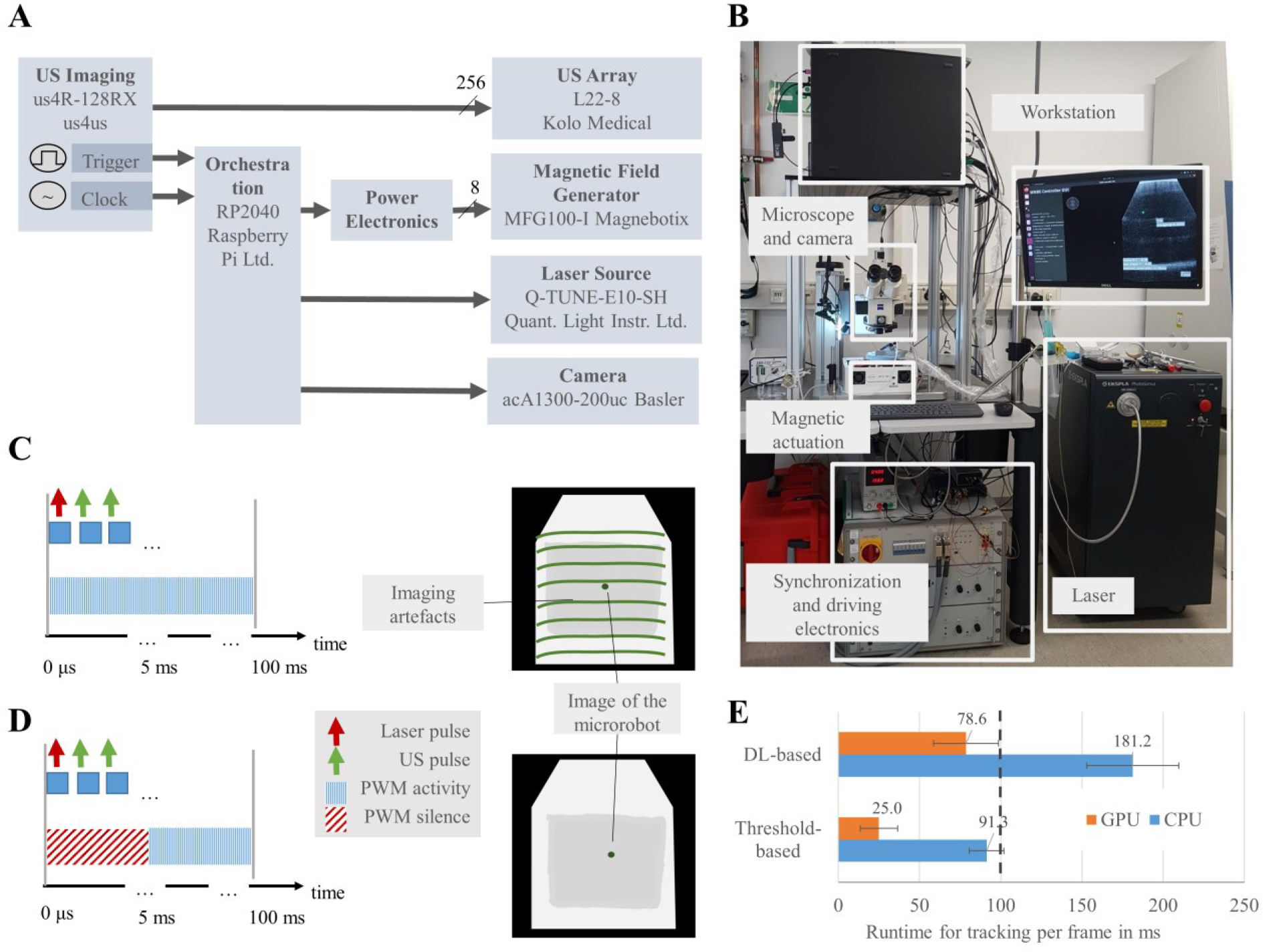
(A) Overview of the system for PA-based closed loop control. The US imaging platform provides a clock signal from which all other timings are derived with a microcontroller-based orchestration module. (B) Photograph of the system with laser safety curtains removed. (C) Timing chart on the left side: Within each imaging period of 100 ms, first the US data acquisition is triggered, followed by the laser pulse and multiple plane wave US emissions afterwards. Right side: Concept drawing of the strong artifacts are observed in the PA image through electromagnetic interference, if the imaging and the magnetic actuation overlaps. (C) Timing chart and concept drawing if the magnetic actuation follows a silence period of 5 ms: The strong artifacts from electromagnetic crosstalk are suppressed. (D) Real-time closed-loop control: The overall timing budget for 10 Hz repetition rate is 100 ms. The computational costs of DL and threshold-based tracking on CPU and GPU are compared.

### B. Synchronization of Imaging and Actuation

Precise timing is crucial for the performance of a PA-based closed-loop control system in two ways: First, beamforming and further processing of a PA image requires low-jitter synchronization between laser emission and US data acquisition. Second, imaging artifacts can be suppressed through synchronization and time-multiplexing. US and PA artifacts stemming from the electromagnetic coupling of the magnetic actuation into the ultrasound signals, as well as overexposure in the auxiliary camera images through the laser pulses can be avoided by sufficient temporal separation. The orchestration of these components is performed by a microcontroller with customized firmware. To prevent having multiple drifting clock sources, the timing is derived solely from the 65 MHz clock of the US platform (Fig. 2).

### C. Deep-Learning based tracking

The detection of microrobots in PA images is challenging, especially in a medical context, because of an often low SNR and an uneven background. DL-based detection may learn to separate the relevant objects from the background more robustly than simple threshold-based methods. “You only look once” (YOLO) version 8 is a model for real-time segmentation and object detection [6]. Its architecture (Fig. 3 A) features the backbone, a structure of descending convolutional layers that reduce the resolution of the input image, a neck that aims to extract contextual information for generalization, and a head section, which generates the bounding boxes [7]. For this work, we selected the “small” model variant, which has 11.2 million parameters and requires 28.6 billion floating point operations when operating on 640 pixel wide images.

**Figure 3:**
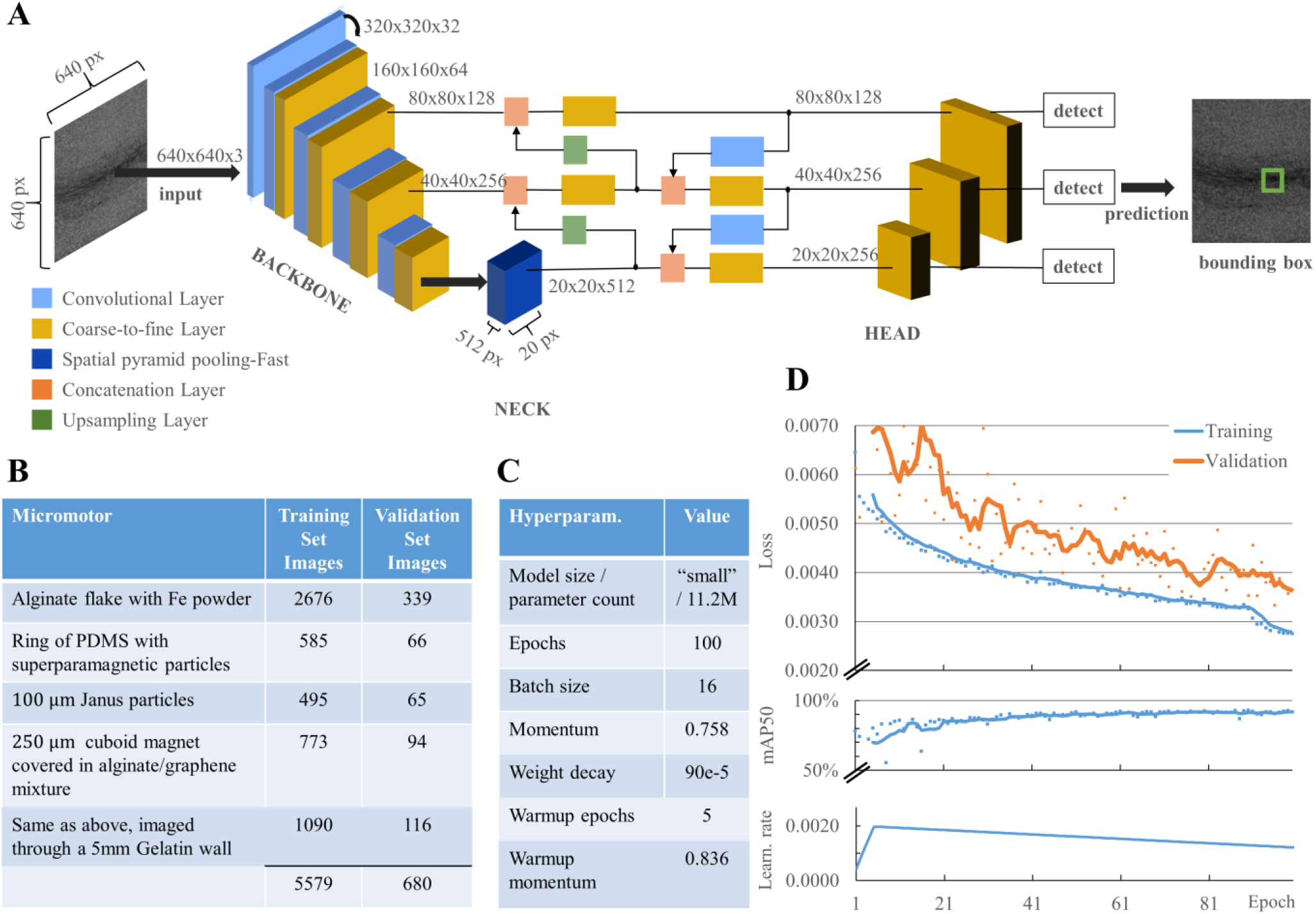
Deep Learning based detection of Microrobots; (A) The architecture of the “You only look once” (YOLO) v8 small model[6] for localizing microrobots. Diagram adapted from [7]. (B) Custom dataset obtained from a diverse spectrum of microrobots. Training images are used for fine-tuning the model and validation images to evaluate the model on unseen data. Note that the spiral-shaped microrobot is excluded from the dataset to test the model’s generalization capabilities. 38% of the images are obtained by augmentation. (C) Hyperparameters and (D) progression of loss [6], mean average precision (mAP50) and learning rate for fine-tuning the pre-trained model on the task of microrobotic localization over 100 epochs.

To fine-tune the pre-trained model on the task of microrobotic localization, we created a custom dataset (Fig. 3 B) for supervised learning. A selection of one-fifth of the frames from long-duration closed-loop experiments is used for either training (5579), validation (680), or testing (724). The test set of images is not used in this study but retained for evaluation after Hyperparameter tuning in the future. The ground truth for the predictions is obtained from threshold-based tracking. of the logarithmic PA amplitude from a diverse spectrum of microrobots in different locations. The spiral-shaped microrobot used in the evaluation is not present in the dataset, to test the model’s capabilities to generalize. 38% of the images of the dataset are obtained by augmentation by computing the complex-valued sum of images with microrobots in different locations. Half of the augmented data is an overlay of two images, the other half of three images. The training was performed over 100 epochs, which reduced the loss in the validation dataset (consisting of 680 images) by approx. 50% (Fig. 3 C and D).

### D. Experimental Setup for Characterization

For demonstrating and characterizing the closed-loop control system, a 3d printed vessel filled with water holds the US probe (L22-8, Kolo Medical) forming an imaging plane over a glass slide at the center of the magnetic actuation system. The laser is guided to the imaging plane through a diffuse reflector. Simultaneous, optical imaging from the top is be performed through a microscope (Zeiss, Oberkochen, Germany) and a camera (acA1300-200uc, Basler AG, Germany).

A 300 μm diameter, spiral-shaped microrobot, which was fabricated using two-photon polymerization of negative tone photoresist (Photonic Professional GT 3D, Nanoscribe GmbH, Germany) as described by Schwarz et al. [9] is exposed to a rotating magnetic field (B = 25 mT, f_rot_ = 0.5 Hz). Due to a ferromagnetic coating, a torque is exerted so it follows the rotation of the field. If in contact with a surface, an out-of-plane rotation will induce a rolling motion whose direction is determined through the direction of the rotational plane. An overview of the parameters and components of the experimental setup is given in Table 1.

**Table 1:** Components and parameters of the experimental setup.

|  |  |
| --- | --- |
| Micro-object | 300 $\mu\text{m}$ spiral-shaped microrobot[9] coated with Fe and Au |
| US probe | 256 elements CMUT phased array (L22-8 Kolo Medical), pitch: 0.2 mm, central frequency: 17 MHz, 77% bandwidth at -6 dB |
| US sequences | Multiplexed PA acquisition and plane wave 5 angles, single cycle centered at 16 MHz, 30 V, $\lambda = 90.6 \mu\text{m}$, 10 frames per second, pulse repetition interval 180 $\mu\text{s}$, TGC gain= 25dB + 150 dB/m, |
| Laser excitation | Q-TUNE-E10-SH, Quantum Light Instruments Ltd., Lithuania, rep. rate 10 Hz, wavelength 1053 nm, pulse length 3.63 ns, pulse energy: 54.32 mJ |
| Actuation | MagnebotiX MFG100i with field enhancement module 10 mm, $\ B\ = 25 \text{ mT}$ , $f_{\text{rot}} = 0.5 \text{ Hz}$ |

## III. Results and Discussion

The rotating magnetic field makes the microrobot roll at the bottom of a vessel and induces a forward motion. Through closed-loop control, an approx. 1.6 mm wide figure-of-8 shape could be followed at a distance of 20 mm from the transducer (Fig. 4 A). Tracing the figure-of-8 shape multiple times over the course of 130 mm gives an average velocity of the microrobot of (467.4 ± 114.4) μm/s.

**Figure 4:**
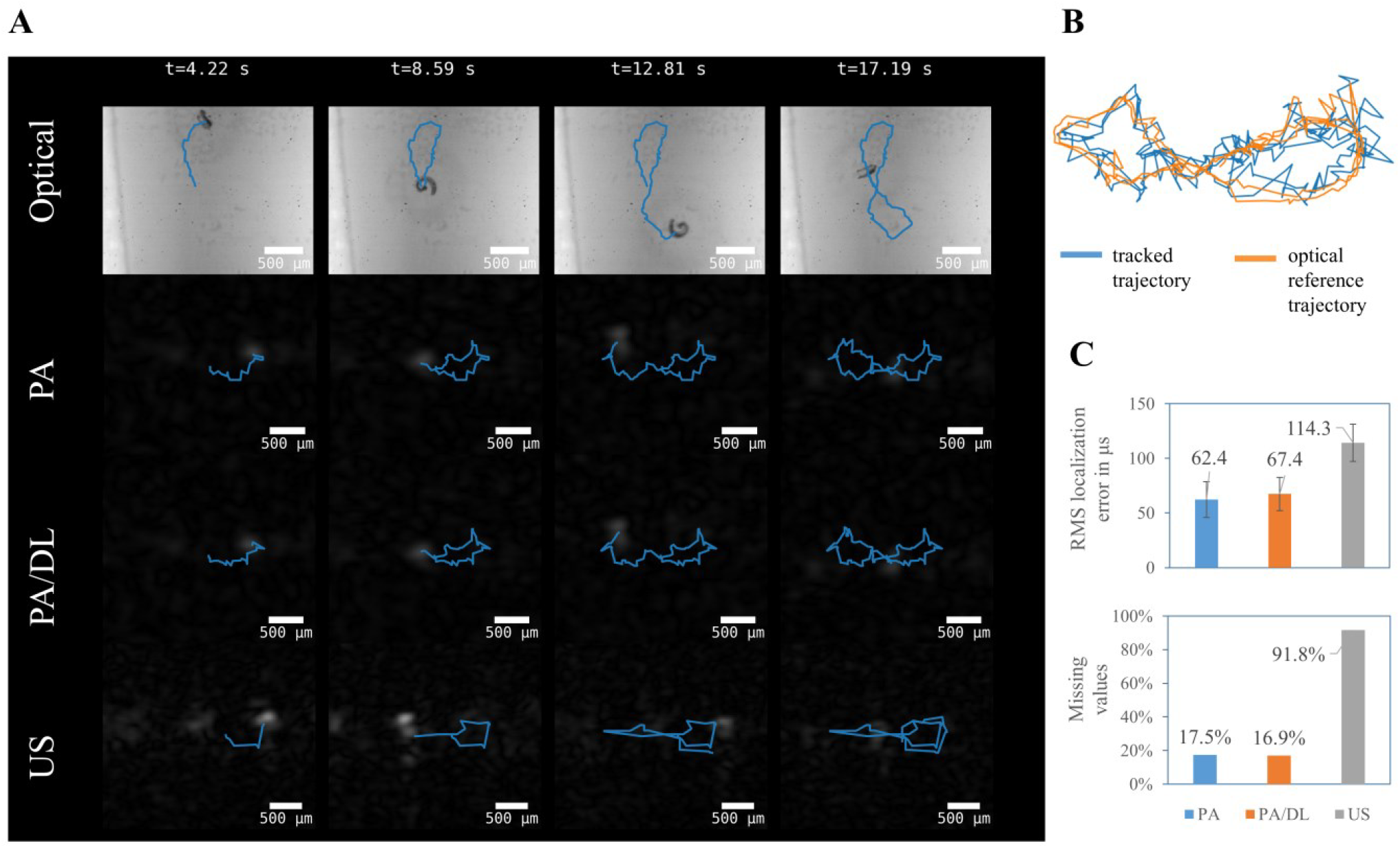
Extract of a dataset for characterizing the tracking performance over a total traveled distance of 130 mm: A 300 μm spiral-shaped microrobot is following a figure-of-8 shape in approx. 20 mm distance of the transducer, with the experimental parameters given in Tab. 1 (A) Trajectories (blue, median filtered over 500 ms) obtained through tracking from optical, PA and US images. (B) Overlay of PA-tracked trajectory and affine transformation of the optical reference over a tenth of the total travel length. (C) Comparison of the root mean square (RMS) localization error and missing detections of the different modalities and tracking methods.

The localization error of the different tracking methods is determined against optical tracking as the gold standard: The trajectory obtained via threshold-based tracking on the camera image is aligned via an affine transform to provide a ground truth (Fig. 4 B). Then the root-mean-squared (RMS) localization error and the percentage of missed detections is calculated (Fig. 4 C). A threshold-based localization on the US image gives more than 91% missed detections, while the remaining points have a localization error of (114.3 ± 17.1) μm. Using threshold-based localization on PA images reduces the missed detection rate to 17.4% and the RMS detection error to (62.4 ± 16.4) μm. By using DL-based tracking, the missed detection rate can be further reduced to 16.9%, while the localization error increases slightly to (67.4 ± 15.3) μm. This demonstrates the robustness of the DL-based tracking model and its ability to generalize, as the microrobot used was not present in the training dataset.

## IV. Conclusion and Outlook

We designed a flexible system for closed-loop control of magnetic microrobots through dual-mode US and PA imaging. It is based on commercially available components, especially an open US platform and a magnetic actuation system, in combination with custom orchestration and driver electronics. Special attention is paid to the synchronization of imaging and actuation, which enables their coexistence without imaging artifacts.

We demonstrated real-time closed-loop control of a 300 μm spiral-shaped microrobot with PA-based feedback. We characterized the localization uncertainty of the system through comparison with optical tracking data. The PA-based tracking outperforms US tracking and achieves an RMS localization error below one-fourth of the body length of the microrobot.

Furthermore, we implemented real-time DL-based tracking which exhibited similar performance to threshold-based tracking. The evaluation on previously unseen experimental data with a new type of microrobot demonstrates the algorithm’s robustness and its capabilities to generalize.

The presented closed-loop control system aims to provide a platform for research in medical microrobotics, with the goal of operating microrobots in deep tissue. Further research will optimize the model through hyperparameter tuning, characterize the system’s performance with ex vivo tissues or phantoms, and investigate the influence of different and dynamic environmental conditions. Finally, we plan to apply the system to *in vivo* microrobotic experiments.

## V. Acknowledgment

The authors like to thank Thorsten Seidemann and Michel Neubert for designing and building the custom power electronics as well as Cornelia Geringswald for 3d printing parts of the experimental setup.

